# Environmental and particle trait controls on microbial carbon turnover on fast-sinking particles in the Iceland Basin

**DOI:** 10.64898/2026.11.13.766158

**Authors:** Cordelia Roberts, E. Elena García-Martín, Carol Robinson, Sabena Blackbird, Stephanie Henson, Emma Louise Cavan

## Abstract

Microbial remineralisation of sinking particles governs the depth and magnitude of organic carbon sequestration in the ocean. Yet, the combined influence of environmental gradients and particle morphology in regulating remineralisation remains poorly constrained. During spring in the subpolar Icelandic Basin, we estimated microbial carbon turnover, a proxy for remineralization, on fast-sinking particles collected at two depths below the mixed-layer using shipboard incubations. Alongside this we used imagery to determine the morphology and complexity of the particles. Microbial carbon turnover exhibited strong depth-dependent variability, with generally higher rates in shallower samples and substantial variability among stations and sampling days. Fast-sinking particle assemblages were dominated by aggregates and detrital material, while faecal pellets comprised only a minor fraction of particles, and particle diversity decreased with depth. Environmental variables, particularly chlorophyll*-a* concentration, explained over 80% of observed variability in microbial carbon turnover. Particle traits describing size, size heterogeneity and structural complexity were also significant predictors, increasing model explanatory power to 91%. Larger and more heterogeneous particle assemblages were associated with lower microbial carbon turnover, while increased structural complexity was associated with lower turnover after accounting for particle size. Together these findings indicate that particle traits provide additional controls on microbial carbon turnover beyond environmental variables alone.

## 1. Introduction

Understanding the contribution of marine microbial organisms in regulating the fate of carbon is central to quantifying the role the ocean plays in storing and cycling carbon. Photosynthetic microorganisms in surface waters fix dissolved inorganic carbon (DIC), in the form of carbon dioxide (CO_2_) to produce organic carbon as biomass, which can sink as particles below the euphotic zone or can be grazed upon by zooplankton which produce fast sinking faecal pellets. The export of material from the euphotic zone into the mesopelagic is mediated by both physical and biological mechanisms including gravitational sinking of particles, vertical mixing, subduction, and active transport by vertically migrating organisms (Boyd et al., 2019) as well as microbial remineralisation and zooplankton grazing (Buesseler and Boyd, 2009; Le Moigne et al., 2025) However, projections of carbon export remain highly uncertain because biological and physical controls on particle flux are not well constrained (Henson et al., 2022). This includes how microbial remineralisation and particle characteristics influence how much carbon can be exported below the mesopelagic where it may be sequestered.

Microbial remineralisation driven by degradation of particles in part sets the sequestration depth of carbon (Boyd et al., 2024; Petiteau et al., 2025). Microbial remineralisation of particulate organic carbon (POC) to dissolved organic carbon (DOC) releases CO_2_ during respiration (Kong et al., 2021). Closer to the surface, this CO_2_ may be outgassed into the atmosphere on climatically relevant timescales (e.g. decades to centuries), whereas if respiration occurs deeper in the ocean interior, carbon may remain stored in deep water masses and be transported by thermohaline circulation for outgassing on century timescales (Kwon et al., 2009; Baker et al., 2022).

Current understanding of particle-associated remineralisation is largely based on measurements of microbial POC turnover, defined here as the loss or transformation of POC through microbial activity. This has primarily been determined under laboratory conditions, and highlights variability in turnover with temperature, pressure, particle composition, lability, and microbial community structure (Iversen & Ploug, 2013; Enke et al., 2019; Nguyen et al., 2022). Recent *in situ* experiments suggest that microbial remineralisation accounts for 7–29% of POC flux attenuation across contrasting oceanic regimes, with its relative importance varying substantially with depth (Boyd et al., 2024). This variability likely reflects changes in environmental conditions, including temperature, oxygen availability and productivity, as well as differences in particle properties encountered throughout the water column. Although environmental conditions such as temperature and oxygen have been shown to constrain microbial carbon turnover (Iversen and Ploug, 2013; Cavan et al., 2017; Cavan and Boyd, 2018) and same particle types found in different sites demonstrate variability in carbon export (Durkin et al., 2022); the relative contribution of these environmental drivers compared with intrinsic particle characteristics remains unclear.

Contrasting morphological characteristics of major marine particle types, including faecal pellets and phytoplankton aggregates, may influence microbial processing and carbon turnover because they differ in size, porosity, surface area and sinking velocity which govern microbial colonisation and access to organic material (Fowler et al., 1986; Belcher et al., 2016; Durkin et al., 2022). Laboratory studies have established mechanistic links between particle morphology, including surface area-to-volume ratio, porosity and shape, and microbial degradation rates (Iversen and Ploug, 2010, 2013). More recently, modelling studies have suggested that particle surface area may regulate microbial processing by influencing ectoenzyme-mediated solubilisation of POC, with surface roughening potentially further enhancing microbial access to organic substrates (Anderson et al., 2023).

To address these uncertainties, we characterised particle morphology, measured respiration rates and calculated carbon turnover of fast sinking particles collected using marine snow catchers during a research cruise to the Iceland Basin in the North Atlantic Ocean in spring 2024. We hypothesised that variability in microbial carbon turnover would be linked to a combination of environmental conditions, particle diversity and particle morphological characteristics (hereafter particle traits) defined here as size and structural related features of the particle that can be derived from images. Specifically, we tested whether particle traits provide additional explanatory power for microbial carbon turnover beyond that explained by environmental conditions alone, and whether incorporating particle information improves our ability to understand observed variability in microbial POC turnover.

## 2. Methods

### 2.1 Study site and environmental conditions

Sampling took place during research cruise DY180 (22^nd^ May – 27^th^ June 2024) to the central Iceland Basin, in the north Atlantic Ocean aboard RRS *Discovery* (Figure 1A). The cruise was part of the BIO-Carbon programme which aims to quantify the role of biological processes in ocean carbon storage. The timing of the cruise was intended to capture the peak and subsequent export of the phytoplankton spring bloom in the Iceland Basin. The cruise consisted of a series of “superstations” of ∼3 days duration, interspersed with excursions of ∼3 days to target coccolithophore blooms in the region. The present study focused on three superstations (S3-S5), each occupied for approximately three days and which we sampled on the first and third days (Figure 1B). Marine snow catchers were deployed at these “superstations” (referred to as stations from here onwards) to collect sinking particles for incubations and imaging characterisation.

**Figure 1.**
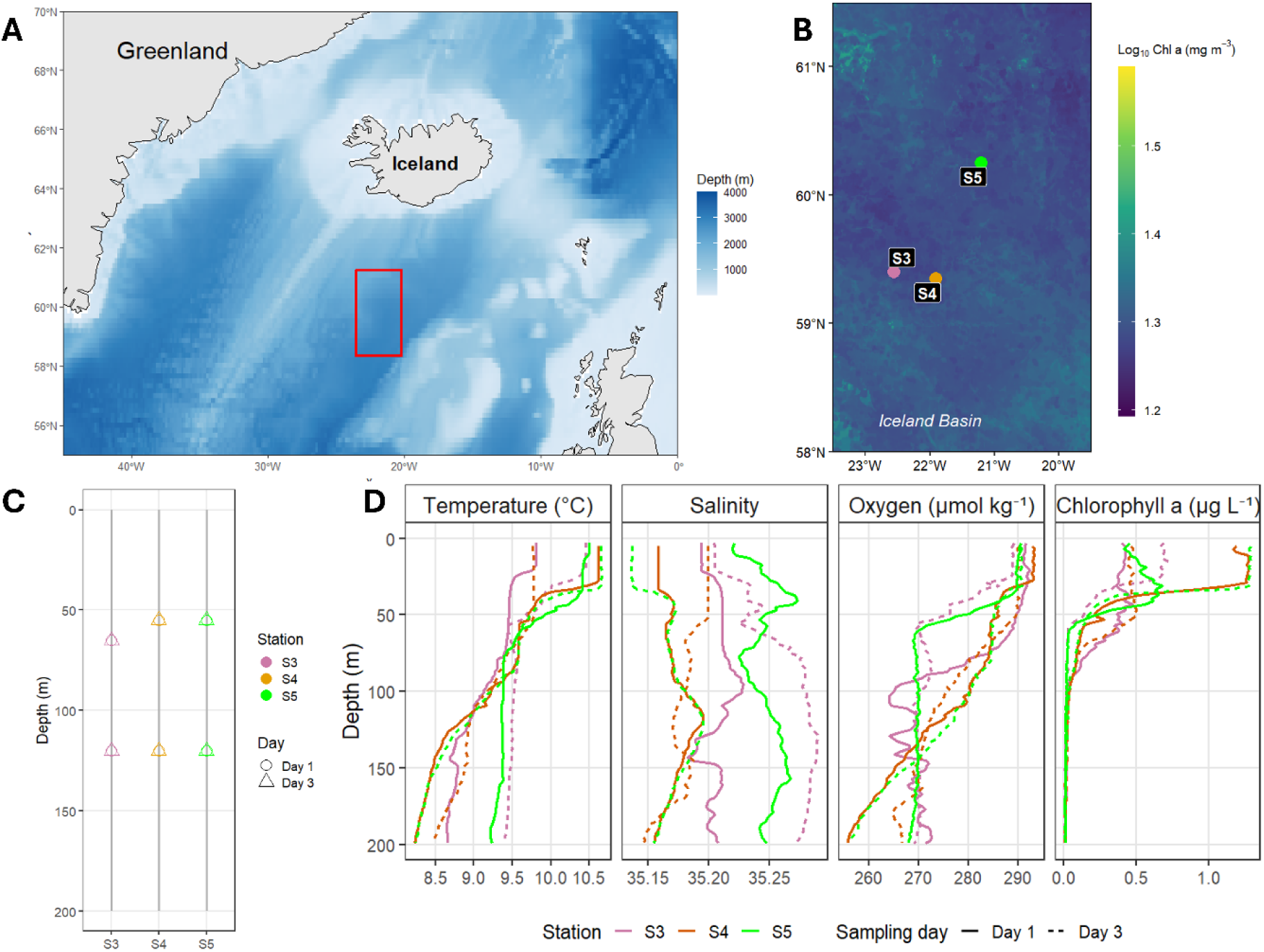
Study region and hydrographic conditions in the subpolar Iceland Basin during May-June 2024. (A) Location of the study area in the North Atlantic Ocean. The red box indicates the spatial extent of the regional chlorophyll *a* map shown in panel B. (B) Location of stations S3–S5 in the overlaid onto mean monthly satellite-derived chlorophyll *a* concentration for June 2024 from the Copernicus GlobColour Ocean Colour product (https://data.marine.copernicus.eu/product/OCEANCOLOUR_GLO_BGC_L4_MY_009_104/description). (C) Vertical profiles of sampling depths. Circles and triangles denote MSC sampling depths on Day 1 and Day 3, respectively. (D) Vertical profiles of temperature, salinity, dissolved oxygen, and chlorophyll *a* from CTD casts conducted two to three hours prior to Marine Snow Catcher (MSC) deployments at stations S3-S5. Colours indicate sampling stations (S3 in pink, S4 in orange and S5 in green), while solid and dashed lines represent Day 1 and Day 3 sampling, respectively.

A conductivity-temperature-depth (CTD) rosette with 24 x 20 L Niskin water samplers was deployed to determine physical characteristics of the water column. Water temperature, conductivity, pressure and dissolved oxygen were measured at each sampling station using a Sea-Bird SBE 9plus CTD fitted with SBE 3P temperature sensors, SBE 4C conductivity sensors, a Paroscientific Digiquartz pressure sensor, Sea-Bird SBE 43 dissolved oxygen sensor, a Chelsea Technologies Group Aquatracka MKIII fluorometer. Oxygen sensors were calibrated against discrete bottle samples collected during the cruise using Winkler titrations.

Mixed-layer depth (MLD) was calculated using a density threshold criterion and defined as the first depth at which potential density exceeded the surface density by 0.05 kg m□³ (Δσ > 0.05 kg m□³; Poulton et al, 2010). For daytime CTD casts, the euphotic depth was calculated from profiles of photosynthetically available radiation (PAR) following Poulton et al. (2010) and Brewin et al. (2015). For CTD casts conducted during darkness, the euphotic zone was estimated as the depth of maximum chlorophyll fluorescence.

Total chlorophyll*-a* was determined by filtering 100 to 250 mL of seawater collected from the Niskin bottles 55 and 120 m onto glass fibre filters (GF/F), with filters placed in 20 mL glass vials and chlorophyll*-a* extracted in 6 mL of 90% acetone for 18 to 24 hours at 4°C in a dark fridge. Chlorophyll*-a* fluorescence was measured using a fluorometer (Turner Designs) with a non-acidification module. The fluorometer was calibrated against a pure chlorophyll*-a* extract prior to the cruise and a solid standard was used at the start of each set of measurements to check for instrument drift. These values were used within the models

### 2.2 Marine snow catcher deployments and particle collection

Marine snow catchers of ∼100 L capacity (MSC, Riley et al., 2012) were deployed at the three stations (Figure 1B), at shallow (55 or 65 m) and deep (120 m) depths on day 1 and day 3 of the 3-day occupation of each station (Figure 1C, Supplementary Table 1). MSC were deployed two-three hours after the CTD deployment beneath the euphotic zone (depth range 15-43m; Supplementary Table 1), which was below or in the lower limit of the MLD on all but one occasion (S3, Day 3). Physical (temperature, salinity, MLD) and environmental (chlorophyll-*a*) characteristics used throughout correspond to the CTD deployment nearest in time before MSC sampling.

After deployment, MSCs were secured on deck for two hours to enable the fast-sinking particles to settle out (Riley et al., 2012). After two hours, the MSC was carefully drained and the bottom tray into which the fast-sinking particles were collected was covered with a lid and opaque black bag to prevent light exposure to samples and removed from the base of the MSC. The tray was transported to a controlled temperature room for subsampling for incubations and particle characterisation. The controlled temperature room was set at *in situ* temperature at ∼ 9.5 °C and under red light for both shallow and deep samples.

### 2.3 Microbial respiration incubations

To estimate the microbial respiration of the free-living and particle attached organisms, subsamples of seawater + particles were collected from the MSC tray into 25 mL glass vials containing PSt3 spots (PreSens, Germany) mounted with silicone glue inside. Three to four replicates were collected from the tray. Microbial respiration was determined by measuring the decrease in dissolved oxygen concentration using Fibox4 OXY10 SMA PreSens optodes (PreSens, Germany). The ten sensors of the OXY10 were two-point calibrated with air-saturated and anoxic waters (precision of ± 0.4% O□ at 100% oxygen saturation). The anoxic water was obtained by dissolving small quantities of sodium sulphite anhydrous in MilliQ water. An initial dissolved oxygen reading was taken after a 30-minute acclimation period in the incubator kept at the respective *in situ* temperatures of the shallow and deep sampling depths as determined by the nearest CTD. Dissolved oxygen concentrations were measured every few hours until the end of the 40-hour incubation period. The incubation duration was selected to ensure sufficient oxygen drawdown for robust estimation of respiration rates while maintaining oxic conditions (>80 % saturation) in all samples.

Prior to measurements, vials were manually inverted in a gentle manner to avoid the formation of oxygen gradients within the vial. Alongside, a control of MilliQ water (MQ) was incubated in the same manner to account for any changes in temperature which could lead to abiotic changes in oxygen during the incubation. Incubations were run at each of the two depths on both day 1 and day 3 at each sampling station. At the end of the incubations, the contents of the vials from the same depth were pooled and filtered onto one pre-combusted (5 hours, 450 °C) 0.7 µm GF/F filter for determination of POC concentration. The filters were stored at –80 °C until analysis. Filtered samples were then dried at 65 °C in an oven for 48 hours and underwent acid vapour treatment with hydrochloric acid for 12 hours to remove inorganic carbon. This was followed by drying at 65 °C for 4 hours prior to analysis.

Organic carbon was determined on four individual quarter pieces from the same filter using a FlashSmart™ elemental analyser (ThermoFisher). A four-point calibration was performed daily using varying masses of High Organic Sediment Standard and Low Organic Soil Standard (Elemental Microanalysis Ltd.; NIST-certified values). Calibration standards were analysed as unknowns at the beginning, middle, and end of each analytical run to assess analytical precision. Carbon mass for each quarter-filter was calculated from the measured carbon content and the dry mass of the respective quarter filter. Whole-filter POC was reconstructed by summing individual quarter-filter measurements.

### 2.4 Microbial particulate organic carbon turnover

To calculate POC-specific respiration or microbial carbon turnover, first we determined the rates of microbial respiration by calculating the slope of the linear regression of dissolved oxygen concentrations (µmol O_2_ L^-1^) over time (h) and assuming a respiratory quotient of 1, following previous particulate respiration studies (Belcher, et al., 2016; Bach et al., 2019; García-Martín et al., 2021). Calculated slopes of the MilliQ vials (0.051 ± 0.046 µmol O□L□¹ h□¹) were subtracted from ‘treatment’ vials to account for any instrument drift, temperature-related variability and abiotic changes in oxygen concentrations. POC associated with each incubation vial was estimated from the POC measured per filter. This was apportioned amongst the incubation vials by dividing by the number of vials filtered on the individual filter and converted to an equivalent concentration (µg C L□¹). POC concentrations were then converted to molar units (µmol C L□¹) using a carbon molar mass of 12.01 g/mol. Microbial POC turnover rates were calculated using Equation 1:

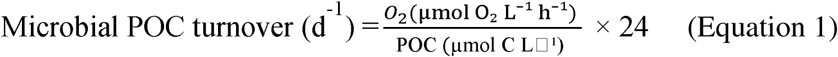

where O□ is the oxygen consumption rate (µmol O□L□¹ h□¹). Oxygen consumption rates were converted to carbon remineralisation rates (µmol C L□¹ h□¹) assuming a respiratory quotient (RQ) of 1, such that 1 mol O□ consumed was equivalent to 1 mol C remineralised. POC is particulate organic carbon concentration (µmol C L□¹) and 24 converts h^-1^ to d^-1^.

### 2.5 Particle imaging, classification and taxonomy

Particle imaging was conducted using FlowCam imaging capabilities. A 12 ml sample of fast sinking material from the sinking tray was collected and fixed with acetic lugol solution (0.3 mL final concentration 2-3%). Samples were stored in 30 mL amber glass bottles in darkness at 4 °C until analysis (∼12 months later). Before analysis, the bottles were inverted gently, and 5 mL of fast sinking material was subsampled. Each subsample was analysed using a FlowCam 8000 series (Fluid Imaging Technologies) fitted with a 5 mL syringe pump. The instrument was operated using an FOV300 flow cell at 4× magnification, with a flow rate of 0.8 mL min□¹ and an efficiency rate of 69.6%. Images were acquired using Auto-Image Mode 7. An equivalent spherical diameter (ESD) acquisition filter was applied, with a minimum particle diameter of 20 μm and a maximum particle diameter of 1000 μm. The FlowCam raw images were exported and processed with Zooprocess software (Gorsky et al., 2010), before being imported into the Ecotaxa web platform (http://ecotaxa.obs-vlfr.fr) for classification. A custom learning set was created from a subset of images which were manually classified, and remaining images were predicted using the EcoTaxa deep learning prediction function (Picheral et al., 2017.) Predicted images were then validated manually. Any images which were classified as blurry, ghost, or artefacts were removed before exporting from EcoTaxa for analysis. Image classifications were used to describe particle assemblage composition and diversity of observed particle types. Particles were grouped into broad categories including detritus, detrital fibres, aggregates, faecal pellets, diatoms, dinoflagellates, ciliates, *Phaeocystis*, copepods, pteropods, radiolaria and unidentified cells. Detritus fibres were classified as elongated, fibre-like detrital material, whereas detritus comprised irregular non-living particulate material that did not exhibit a distinct fibrous morphology. Aggregates were accumulations or aggregations of cells and/or detritus, fluffy in texture and potentially containing interstitial spaces with a non-distinct irregular shape or undulate margin. Morphological descriptors (particle traits) exported from EcoTaxa included equivalent spherical diameter (ESD), aspect ratio, elongation and fractal dimension. Summary statistics describing particle size structure were calculated for each sample, including mean and median ESD, size heterogeneity (coefficient of variation of particle area), and the relative abundance of particles within predefined projected area size classes: very small (<500 μm²), small (500-2000 μm²), medium (2000-5000 μm²), and large (>5000 μm²).

### 2.6 Statistical Analyses

Particle type diversity was quantified using Shannon’s diversity index (H’), calculated from the relative abundance of EcoTaxa particle classification categories within each sample. Bray-Curtis dissimilarities were calculated using the *vegdist* function in the R package vegan (Oksanen et al., 2026). Patterns in assemblage composition were visualised using non-metric multidimensional scaling (NMDS), and differences among stations and depths were tested using permutational multivariate analysis of variance (PERMANOVA).

Differences in microbial carbon turnover between sampling depths were assessed using Wilcoxon rank-sum tests. The effects of depth and station on microbial carbon turnover were further evaluated using linear models that included depth, station and their interaction terms. Relationships between microbial POC turnover and environmental variables were examined using an ordinary least squares linear regression on non-transformed data. Environmental variables considered included temperature, oxygen, salinity, mixed-layer depth and chlorophyll*-a* concentration. Following exploratory analyses and assessment of collinearity, non-transformed chlorophyll*-a*, dissolved oxygen concentration and mixed-layer depth were retained in the final environmental model. A second particle trait-informed model incorporated the PCA-derived trait axes (PC1 and PC2; see Section 3.3) and particle diversity in addition to the environmental variables. Particle diversity was not a significant explanatory variable to explain microbial carbon turnover and therefore not retained in the final particle trait-informed model. Model performance was evaluated using coefficient of determination (R²), Akaike Information Criterion (AIC), and root mean square error (RMSE). As including individual traits directly in regression models can lead to collinearity and over-fitting, a principal component analysis (PCA) was used to derive composite trait axes. A PCA was performed on particle morphological descriptors including equivalent spherical diameter (ESD), object area, Feret diameter, aspect ratio, elongation, fractal dimension, size heterogeneity, and the relative abundance of particle size classes. Prior to analysis, variables were centred and standardised to unit variance to ensure equal weighting of traits measured on different scales.

## 3 Results

### 3.1 Environmental conditions

In the subpolar Icelandic Basin, our observations captured temporal progression in upper-ocean hydrography and biological production (Figure 1D). Temperature, salinity and oxygen profiles revealed vertical and temporal variability in water column structure between and within stations (Figure 1D). The mixed-layer depth (MLD) changed between and within stations (Supplementary Table 1) with a notable deepening event observed at S3 between day 1 (MLD = 25 m) and day 3 (MLD = 78 m). For S3 Day 3, MLD varied between two consecutive CTDs decreasing from 78 m pre MSC deployment to 64 m post MSC deployment. The earlier value was used because it most closely represented conditions at the time of particle collection. In contrast, mixed-layer depth remained relatively stable at S4 (29–37 m) and S5 (33–35 m) between pre and post MSC deployments.

Chlorophyll*-a* concentration also varied among stations and with depth (Figure 1D). At the MSC sampling depths (55–65 m), concentrations ranged from 0.10 to 0.82 µg L□¹, with the highest concentration observed at S3 Day 1 and the lowest at S4 Day 1. Chlorophyll*-a* declined with depth across all stations. However, following mixed-layer deepening at S3, chlorophyll*-a* concentrations at 120 m increased from 0.05 µg L□¹ on Day 1 to 0.22 µg L□¹ on Day 3, resulting in a weaker vertical gradient in chlorophyll*-a* than observed at the other stations (Figure 1D).

### 3.2 Microbial Carbon Turnover

Microbial carbon turnover rates ranged from 0.009–1.95 d□¹ (mean ± SD = 0.49 ± 0.62 d□¹, n = 31) with the highest rates observed at S3 day 1 and the lowest at S5 day 3 (Figure 2). Overall, microbial carbon turnover was significantly higher in shallow than deep samples (Wilcoxon rank-sum test, W = 99, p <0.001), although the depth response differed across stations (depth x station interaction p <0.001). However, S5 exhibited the opposite pattern with higher turnover in deeper than shallow samples on both sampling days (Figure 2). The magnitude of the decline in microbial carbon turnover with depth varied across stations and sampling days, ranging from weak attenuation at S3 Day 3 (0.067 to 0.013 d□¹) to more than an order of magnitude decline from shallow to deep samples at S4 Day 3 (0.86 to 0.06 d□¹; Figure 2). Environmental variables explained 80% of the variance in microbial carbon turnover (adjusted R² = 0.80), including chlorophyll-*a*, dissolved oxygen and mixed layer depth. Turnover was positively associated with chlorophyll-*a* (p <0.001) and negatively associated with dissolved oxygen and mixed-layer depth (p <0.001).

**Figure 2.**
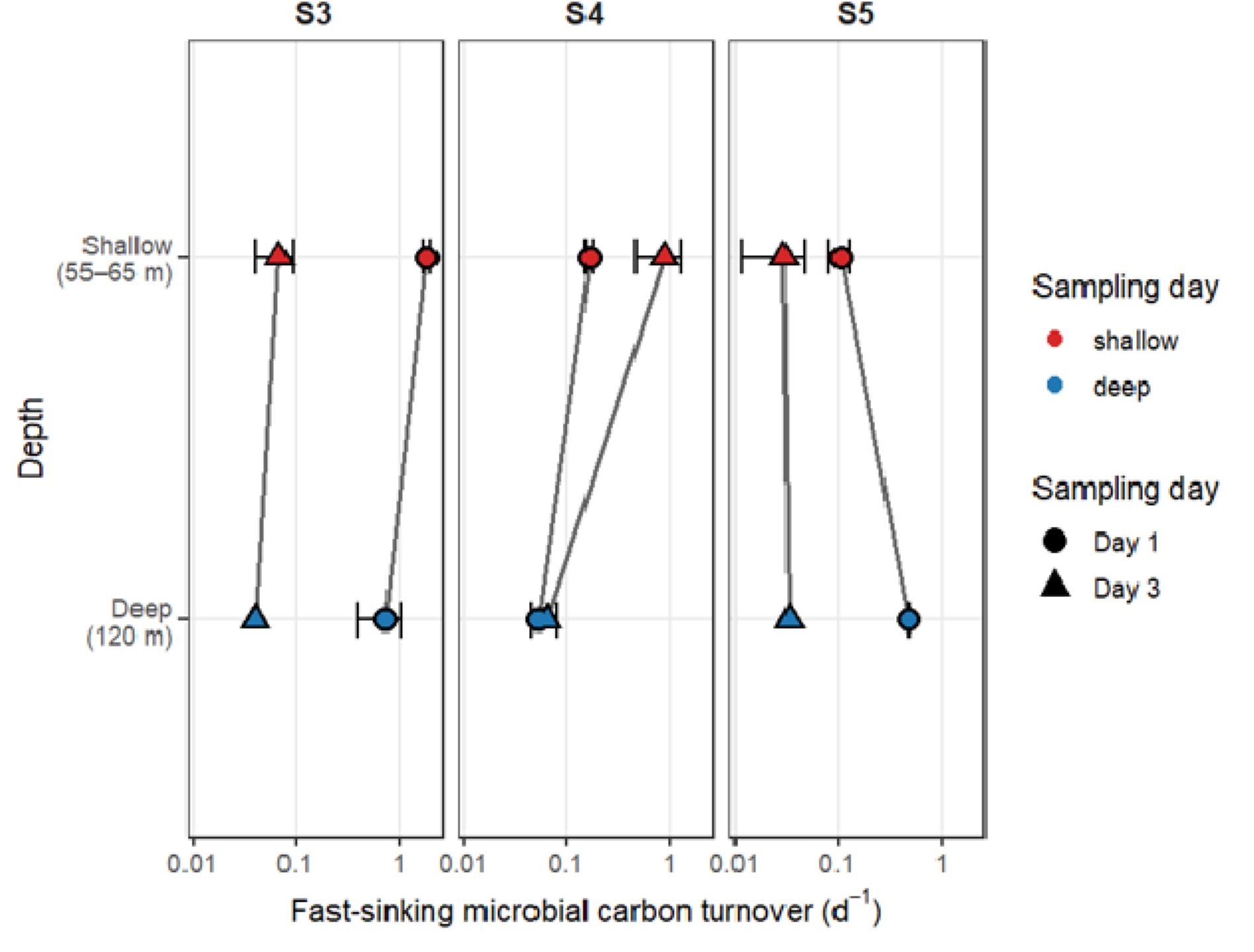
Microbial carbon turnover (d^-1^) associated with POC of fast-sinking particles collected in the Iceland Basin at shallow (red) and deep (blue) depths at stations S3, S4, S5. Symbol represents day 1 (circle) and day 3 (triangle) of station sampling. Error bars represent ±1 SD of replicate incubations (n=3-4). Where no error bars are visible, only a single non-zero turnover estimate remained following temperature corrections, meaning standard deviation could not be calculated.

### 3.3 Particle Diversity and Morphology

Detritus was the dominant particle type across all stations and depths, accounting for on average ∼50% of particles in shallow samples and 65% at depth (Figure 3A). Faecal pellets had a minor contribution to the abundance of fast-sinking particles (less than 1%), while detrital fibres and aggregates were also present in low relative abundances (both less than 5%). Living categories, mainly diatoms and dinoflagellates, comprised around 5-30% of the particles in the fast-sinking tray. Diatoms were more abundant in shallow samples (∼7%) where chlorophyll-*a* concentrations were highest. The ‘other’ category represented between 10-30% of the relative contribution. Ciliates although present, had a very minor contribution to the relative abundance (<0.5%).

**Figure 3.**
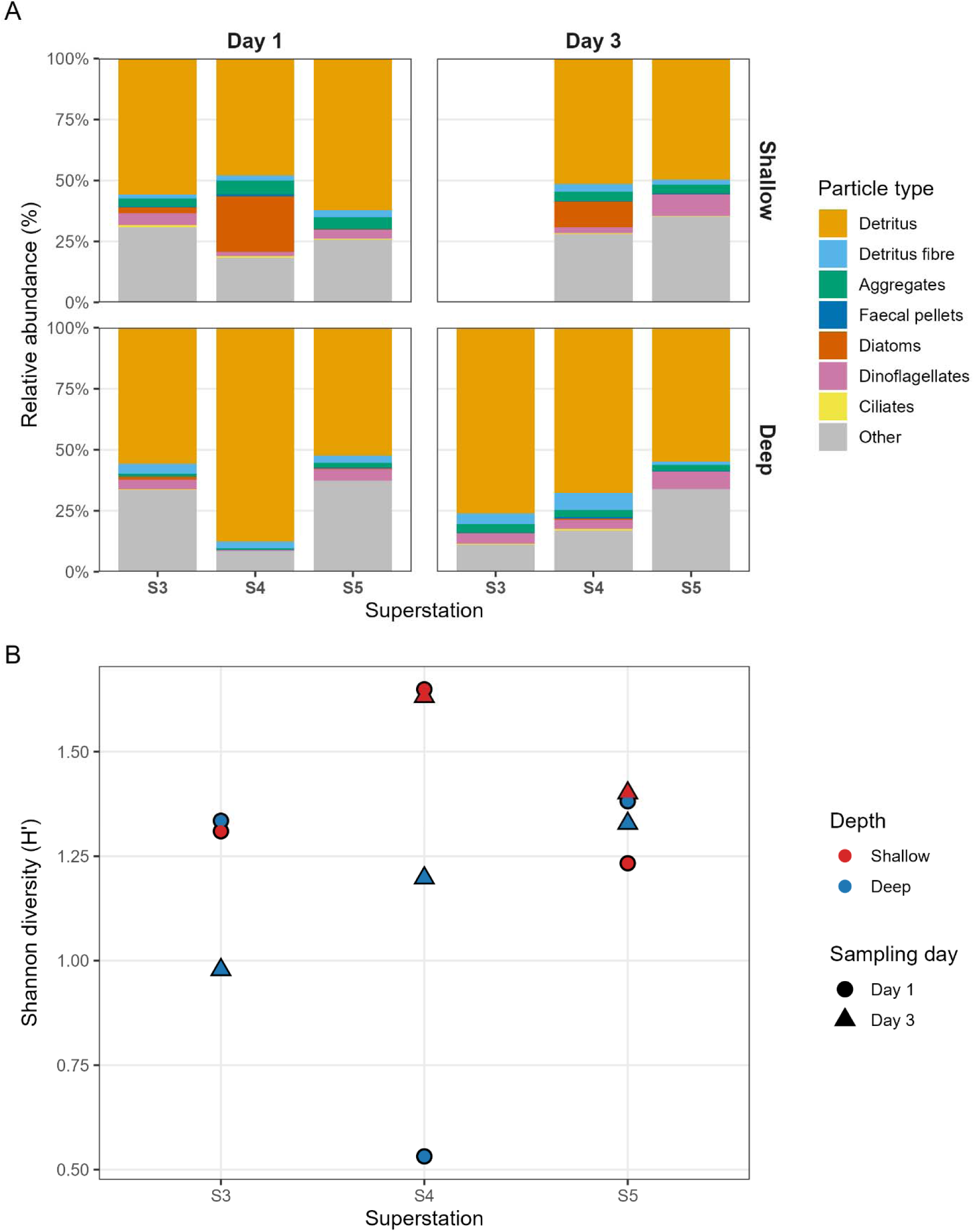
A) Relative abundance and B) Shannon diversity of particle types observed after collection in the fast sinking tray at shallow and deep depths at stations S3, S4, S5. Other includes: *Phaeocystis*, copepods, pteropods, radiolaria and unidentified cells.

Particle assemblage diversity varied across stations and depths. The high abundance of detritus in the deep sample at S4 on Day 1 (Figure 3B) corresponded to a low Shannon’s diversity index. Shallower samples exhibited greater particle diversity than deeper samples (Figure 3B). Bray-Curtis dissimilarity analysis revealed broad overlap in particle type assemblages across stations, with weak clustering across depths and these groupings were not statistically significant (PERMANOVA, p >0.05; Supplementary Figure 1). There was a large variability in morphologies between- and within particle types (Figure 4A-G).

**Figure 4.**
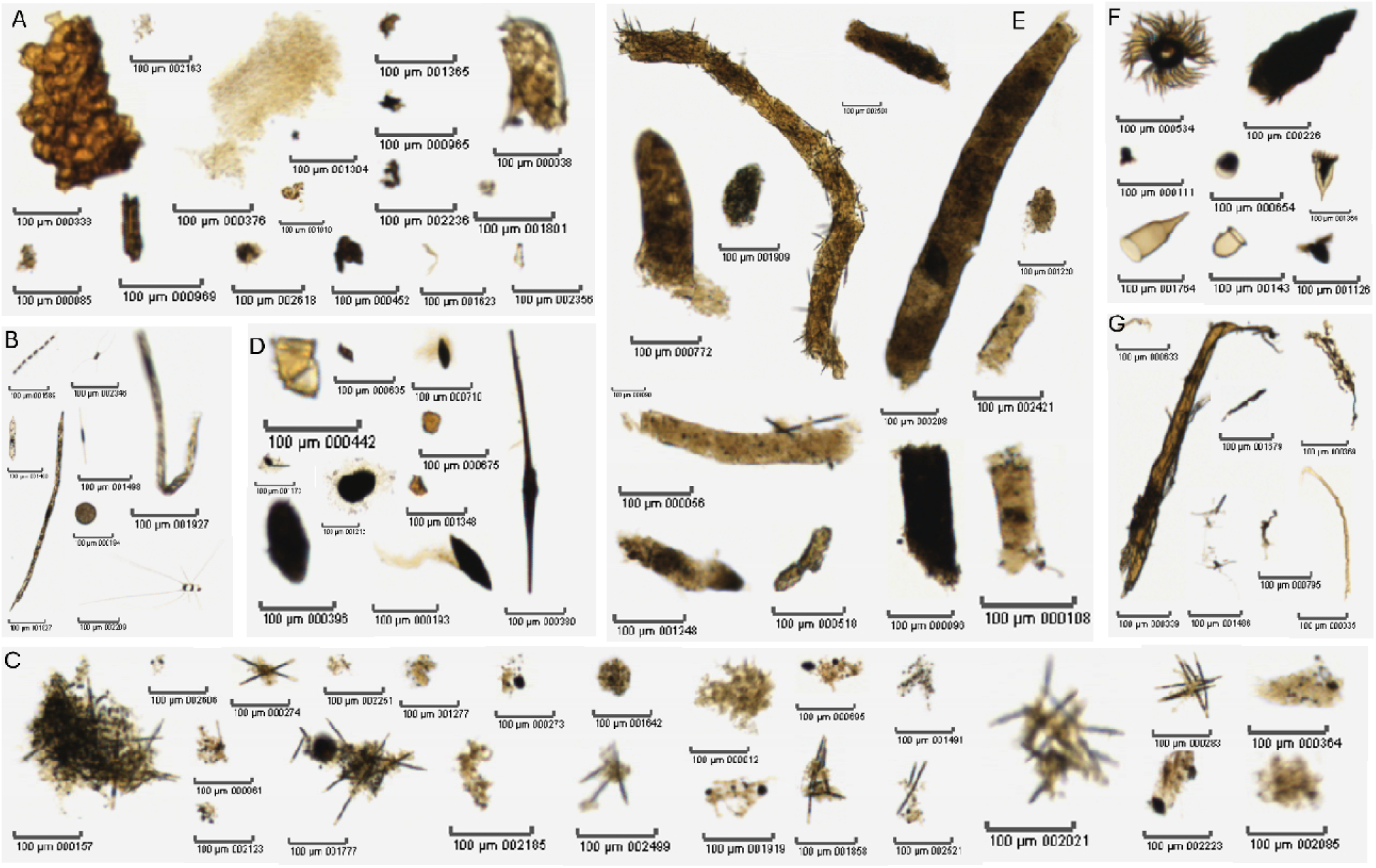
Examples of the variability in morphology of particle types are shown in panels A to G: A) detritus, B) diatoms, C) aggregates, D) dinoflagellates, E) faecal pellets, F) ciliates, G) detritus fibres. Scale bar is 100 µm.

Morphological properties varied among stations, depths and sampling days (Figure 4, Supplementary Figure 2). Mean equivalent spherical diameter (ESD) differed among stations and sampling periods, with the largest particles observed at S4 and the smallest at S5 (Supplementary Figure 2A). Particle shape complexity, represented by fractal dimension, varied over a comparatively narrow range but nevertheless exhibited clear differences at S4 (Supplementary Figure 2B). Size heterogeneity, reflecting variability in particle area within individual samples, showed the greatest differences among samples, with the most heterogeneous particle assemblages observed at S4 (Supplementary Figure 2C). These morphological descriptors were subsequently summarised using principal component analysis into a particle size-related trait axis (PC1) and a structural complexity axis (PC2).

### 3.4 Influence of particle morphology on Carbon Turnover

To investigate whether particle morphological characteristics explained further variance in microbial carbon turnover, we constructed a linear regression model incorporating particle traits identified as key axes of variation in a principal component analysis (Supplementary Figure 3A). Particle traits were treated in a multidimensional space, reflecting the fact that microbial accessibility to carbon likely depends on combinations of particle size, and morphological structure such as edge complexity (fractal dimension) and elongation, rather than on any single characteristic. The first axis (PC1) primarily represented variation in particle size and size-dependent properties, including equivalent spherical diameter (ESD), particle size-class composition and size heterogeneity. The second axis (PC2) captured variation in particle structural complexity and geometry, primarily associated with fractal dimension, elongation, and aspect ratio. The size-related particle trait axis (PC1) showed no significant relationship with chlorophyll*-a* concentration (ρ = -0.21, p = 0.22; Supplementary Figure 3B) but was positively correlated with mixed-layer depth (Spearman’s ρ = 0.51, p = 0.002; Supplementary Figure 3C). Sensitivity analyses excluding the deepest mixed-layer observation yielded a weaker but similar relationship (ρ = 0.46, p = 0.007), indicating that the correlation was not driven by a single influential observation. In contrast, the complexity-related trait axis (PC2) was positively correlated with chlorophyll*-a* concentration (ρ = 0.64, p < 0.001; Supplementary Figure 3D) but showed no significant relationship with mixed-layer depth (ρ = 0.09, p = 0.59; Supplementary Figure 3E).

Our particle trait-informed linear model explained 89% of the variance in fast sinking microbial carbon turnover observed at our study site, improving the model accuracy relative to environmental parameters alone, reducing RMSE from 0.26 to 0.18 and decreasing AIC from 14 to -6. (Supplementary Figure 4). Chlorophyll-*a* was the strongest predictor of microbial carbon turnover (standardised β = 1.04; Figure 4A), followed by dissolved oxygen (β = −0.65; Figure 4A). Particle complexity traits (PC2; β = -0.23) and particle size traits (PC1; β = -0.14) exerted smaller but significant effects, whereas mixed-layer depth was not a significant predictor in the final particle-informed model (β = -0.06, p = 0.29; Figure 5A).

**Figure 5.**
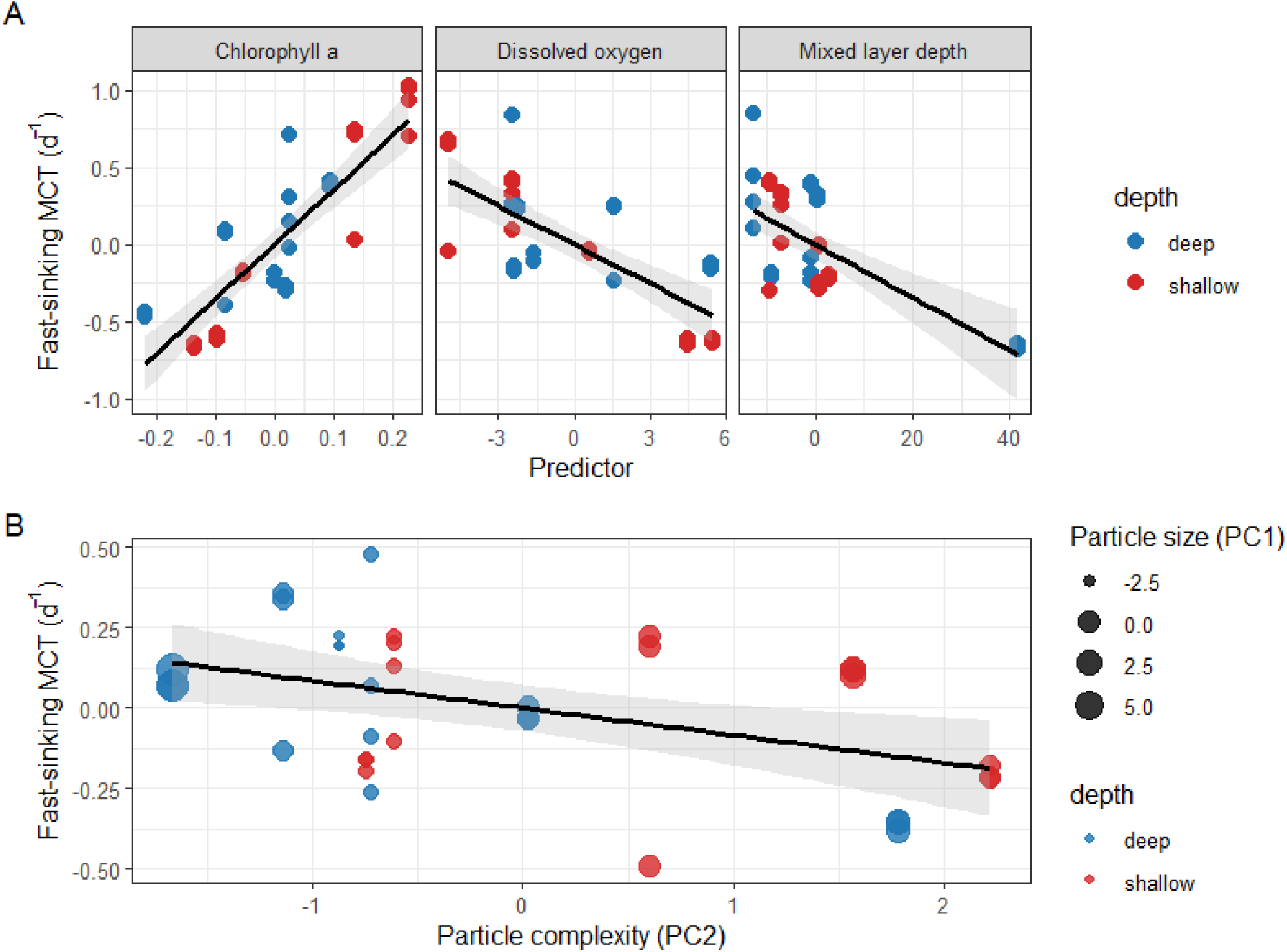
Partial regression plots showing the effect of predictor variables on fast-sinking microbial carbon turnover after accounting for the influence of all remaining predictors in the final model. A) environmental parameters (chlorophyll *a*, dissolved oxygen, mixed-layer depth) and B) particle complexity (PC2) after accounting for the effects of the remaining predictors. Points represent the partial residuals rather than raw observed values. Plotted are non-dimensional values extracted from the regression model. Values above zero indicate microbial carbon turnover that is higher than expected after accounting for the effects of the remaining predictors in the model, whereas values below zero indicate turnover that is lower than expected. Negative values therefore represent residual variation and do not indicate negative microbial carbon turnover rates. Size of the point on panel B represents particle size as a function of the PC1 axis (size) identified in the PCA (Supplementary Figure 3A). Grey shading represents 95% confidence intervals.

Mean equivalent spherical diameter (ESD) and mean fractal dimension were major drivers of the PC1 and PC2 axes respectively, but when considered in isolation showed no relationship with fast sinking microbial carbon turnover (Supplementary Figure 5A, 5B). However, after accounting for covariation amongst environmental and particle traits, both PC1 and PC2 emerged as significant predictors of microbial carbon turnover (Supplementary Table 2). Partial regression analyses showed that turnover declined with increasing particle size-related traits and particle complexity-related traits, indicating particle traits explained additional variance in turnover beyond that associated with chlorophyll-*a*, oxygen concentration and mixed-layer depth (Figure 5B). This suggests that particle assemblages characterised by larger particles and greater structural complexity tended to exhibit lower microbial carbon turnover after accounting for environmental variability.

Fast-sinking particles generally exhibited higher carbon turnover at 55-65 m than at 120 m. However, the magnitude of this shallow-deep difference varied among stations and sampling days. Deeper samples were generally characterised by lower turnover and larger particle size-related trait values, suggesting that variation in particle traits may contribute to depth-dependent patterns in microbial carbon turnover. Particle complexity (PC2), chlorophyll-*a*, and dissolved oxygen exhibited similar relationships with turnover across depths. Although particle diversity varied across stations and depths, it did not explain additional variance in carbon turnover when included in the multivariate model (p > 0.05).

## 4 Discussion

Our field-based observations demonstrate that particle morphology, which is not explicitly represented in most biogeochemical models beyond simple particle size classifications, can improve the explanation of variability in microbial POC turnover when considered alongside environmental variables. Environmental conditions alone explained the majority (80%) of observed variability in remineralisation rates. The inclusion of particle trait information further increased explanatory power, indicating that particle morphology captures additional variability not explained by environmental conditions alone. These findings suggest that environmental context and additional particle traits may represent useful, but currently under-utilised, explanatory descriptors for understanding variability in microbial carbon turnover. However, further studies across a broader range of oceanographic regions and seasons are required to validate our initial relationships.

### 4.1 Environmental controls on microbial POC turnover

Environmental variation explained the largest control on carbon turnover in fast sinking material. Microbial carbon turnover in fast-sinking material was generally higher at shallower depths than deeper depths across stations and sampling days. Higher turnover in shallower samples was accompanied by elevated respiration rates and generally higher concentrations of POC in the incubation vials, indicating more rapid microbial processing of a larger organic matter pool (Supplementary Figure 6). Previous studies have shown that degradation of sinking particles is influenced by substrate quality and the progressive removal of labile compounds during transport through the water column (De Baar et al., 1983; Wakeham et al., 1984; Iversen and Ploug, 2013). Because turnover reflects the ratio of respiration to particulate organic carbon, higher turnover in shallow samples likely reflects differences in both substrate availability and microbial respiration relative to carbon stocks. Although lower temperatures can suppress remineralisation rates (Iversen and Ploug, 2013; Cavan and Boyd, 2018), no relationship was observed between temperature and microbial carbon turnover in this study. This may reflect the relatively narrow temperature range encountered across samples (9-10 °C), which limited our ability to resolve temperature effects.

Microbial carbon turnover declined from our first station (S3) to the final one (S5) in tandem with declining chlorophyll-*a* concentrations. As chlorophyll-*a* emerged as a significant predictor in the environmental model, this result suggests that spatial variability in microbial carbon turnover was linked to variation in surface productivity and organic matter supply across the study region. This interpretation is consistent with previous studies demonstrating that microbial remineralisation is strongly influenced by the quantity and quality of phytoplankton-derived organic matter and the progressive loss of labile compounds during particle transport and degradation (De Baar et al., 1983; Wakeham et al., 1984; Iversen and Ploug, 2013). Mixed-layer depth was a significant predictor in the environmental model and was positively associated with the particle size-related trait axis (PC1). However, mixed-layer depth was not retained as a significant predictor in the final particle-informed model, suggesting that part of its explanatory power may be captured by variation in particle traits. Variability in mixed-layer depth can influence phytoplankton growth, particle production and export dynamics by altering light and nutrient availability, thereby modifying the quantity and characteristics of organic matter available for microbial remineralisation (Giering et al., 2016).

### 4.2 Particle diversity and morphological control on microbial POC turnover

Fast-sinking particles were dominated by detrital material across stations and depths, consistent with previous observations from the Northeast Atlantic (Lampitt et al., 1993, 2001) Nevertheless, fast-sinking particles showed contrasting patterns in diversity between depths. Reduced particle diversity with depth has been interpreted as evidence of selective attenuation of labile particle classes during sinking (Trudnowska et al., 2021) If this mechanism were important within our observations, a relationship between particle diversity and microbial carbon turnover might be expected. The absence of a relationship between particle diversity and microbial carbon turnover suggests that morphological characteristics captured aspects of particle variability that were more relevant to remineralisation than particle identity alone.

Microbial carbon turnover was negatively related to the particle size-related trait axis (PC1). Assemblages characterised by larger and more heterogeneous particles exhibited lower microbial carbon turnover, consistent with evidence that increasing particle size reduces surface-area-to-volume ratios and may limit microbial access to organic substrates (Iversen and Ploug, 2013; Logan and Wilkinson, 1991). Particle size-related traits were positively associated with mixed-layer depth, suggesting that upper-ocean structure influenced particle trait distributions across stations, and remained significant after removal of the deepest mixed-layer observation, indicating it was not solely driven by a single point. The relationship between PC1 and microbial carbon turnover indicates that particle size-related characteristics represent an important control on remineralisation rates beyond environmental variables alone. Because particle morphology is shaped by both particle formation and degradation processes, it may integrate information about particle history that is not captured directly by environmental measurements. Consequently, particle size-related traits may reflect underlying variation in substrate accessibility, particle age and organic matter composition, all of which are known to influence microbial remineralisation (Enke et al., 2018; Iversen and Ploug, 2013; Nguyen et al., 2022).

Beyond particle size, microbial carbon turnover was also influenced by structural complexity. Structural complexity was negatively associated with particle size, with smaller particles generally exhibiting higher fractal dimension values than larger particles (Supplementary Figure 7; ρ= -0.5, p <0.01). This is consistent with enhanced surface-area-to-volume ratios promoting microbial colonisation and accessibility to organic substrates (Logan and Wilkinson, 1991; Ploug and Grossart, 2000). Recent modelling studies suggest microbial degradation can increase particle surface area through structural roughening, thereby enhancing opportunities for enzymatic solubilisation (Anderson et al., 2023). However, after accounting for particle size, (Enke et al., 2018; Hedges et al., 2001) increased structural complexity was associated with lower microbial carbon turnover. This suggests that structural complexity may capture aspects of particle formation and degradation life-history, including aggregation, ageing and physical restructuring, rather than simply reflecting increased microbial accessibility. Similar decoupling between particle structure and microbial carbon processing has been reported previously, where increased structural irregularity associated with particle ageing and physical protection coincided with declining substrate availability (Hedges et al., 2001; Enke et al., 2018).

At our final station (S5), carbon turnover at 120 m exceeded that observed in shallower samples. This pattern reflected higher respiration relative to particulate organic carbon stocks, rather than elevated respiration alone. Although particle characteristics also differed between sampling periods, corresponding changes in turnover were not always observed. This suggests that particle morphology alone does not fully explain variability in microbial carbon turnover and that additional factors, including organic matter composition (Iversen and Ploug, 2013), substrate lability (Nguyen et al., 2022), and the composition and activity of particle-associated microbial communities (Datta et al., 2016) may have contributed to the observed variability in microbial carbon turnover.

### 4.3 Limitations

In some cases, turnover rates exceeded 50% d□¹ (n=9) and 100% d□¹ (n=7) in incubations (total n= 38), indicating rapid carbon cycling. Conversely, several deep samples produced oxygen consumption rates that were similar to, or lower than, those observed in MilliQ controls (n=7). These observations of low oxygen consumption may be approaching the lower analytical limits of the optodes (precision of ± 0.4% O□ at 100% oxygen saturation). For rapid turnover rates, our oxygen consumption measurements may have included contributions from particle-associated metazoans e.g. copepods that remained within incubation vials despite efforts to remove visible swimmers. In addition, dissolved organic carbon was not quantified during the incubations, despite it representing a key component of organic carbon released from particles (Noji et al., 1999; Alldredge, 2000) and it being the substrate directly assimilated by microbes following extracellular solubilisation of particulate material (Arnosti, 2011; Arnosti et al., 2012). As turnover rates were calculated using POC, as per previous studies on fast sinking particles (Cavan et al., 2017; García-Martín et al., 2021), respiration supported by unmeasured DOC could lead to an overestimation of true particulate carbon degradation. Consequently, turnover may reflect processing of both particulate and dissolved carbon pools. This may also contribute to the unexpectedly strong environmental relationships observed, although the relative contribution of dissolved versus particulate substrates cannot be determined from the present study. Additionally, as these are particle-enriched incubations there is likely still a contribution from the free-living microbial respiration which has not been directly quantified. Therefore, our reported turnover rates should be considered an upper estimate of microbial utilisation of particulate carbon. Future studies that simultaneously quantify POC, DOC and respiration are required to determine whether environmental variables exert similarly strong control over particulate organic carbon degradation specifically.

## 5 Conclusion

Together, these findings demonstrate that microbial carbon turnover on fast-sinking particles is structured by depth-dependent changes in environmental conditions and particle characteristics. Turnover rates were consistently higher in shallow samples within the mixed layer than at 120 m, where particle morphological diversity was also highest. Environmental variables explained more than 80% of the observed variation in microbial carbon (POC + DOC) turnover, and the inclusion of particle traits increased model explanatory power to 91%. These results highlight environmental variables as primary predictors of microbial carbon turnover, and particle morphology provide complementary predictive power. We suggest that incorporating particle traits alongside environmental variables may improve predictions of particle remineralisation and the efficiency of biological carbon sequestration in the ocean.

## CRedIT authorship contribution statement

C. Robinson, E.E.G.-M, and E.L.C conceived and designed the study. C. Roberts and E.E.G.-M. conducted field sampling, incubation experiments during cruise DY180 and run fast-sinking particles through the FlowCam. S.B performed particulate organic carbon analyses. C. Roberts analysed the data, interpreted the results with input from all co-authors, prepared the figures, and wrote the first draft of the manuscript. C. Robinson, E.E.G.-M, S.H. and E.L.C. contributed to data interpretation and manuscript development. E.L.C., C. Robinson, E.E.G.-M and S.H. secured funding for the project. All authors reviewed and approved the final manuscript.

## Acknowledgements

We thank the officers, crew and scientists of RRS *Discovery* cruise DY180 for their support during field operations. This work formed part of the Natural Environment Research Council (NERC)-funded BIO-Carbon programme and the PARTITRICS project. We thank Chelsey Baker and Will Major for support in deploying the marine snow catchers and particulate organic carbon sampling. We are grateful for the calibration of CTD sensors by Ed Mawji with discrete dissolved oxygen measurements and thank Alex Poulton and Ben Gustafson for discrete chlorophyll*-a* analyses. We are also grateful to Yvonne Firing for CTD data processing, Amanda Elineau for assisting with ZooScan transfer of FlowCam imaging data and EcoTaxa training, and Sarah Cryer for calculating mixed-layer and euphotic depths. We also extend thanks to the Icelandic Coast Guard for their assistance during the medical evacuation conducted during the cruise.

## Data availability

The datasets generated and analysed during this study are available through the British Oceanographic Data Centre (BODC) at DOI:60154D447B24990DE0637086ABC0ABB9 FlowCam data used in this study can be found on EcoTaxa (https://ecotaxa.obs-vlfr.fr/prj/19681).

