## Supplementary for "Environmental and particle trait controls on microbial carbon turnover on fast-sinking particles in the Iceland Basin"

This file includes:

Supplementary Figures 1-7

Supplementary Tables 1-2

**Supplementary Figures**

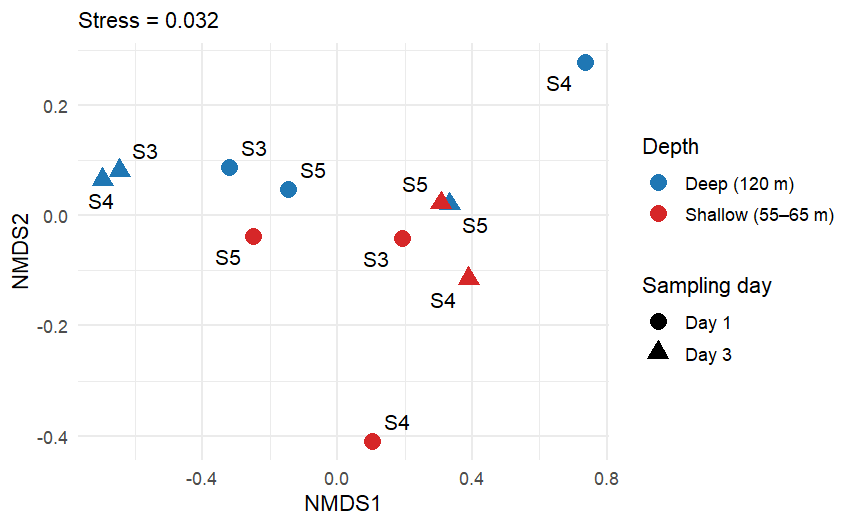
**Supplementary Figure 1.** Non-metric multidimensional scaling (NMDS) plot of normalised fast sinking particle assemblages (relative abundance of particle types) for shallow (red) and deep (blue) depths at stations S3, S4, S5. Symbol represents day 1 (circle) and day 3 (triangle) of station sampling.

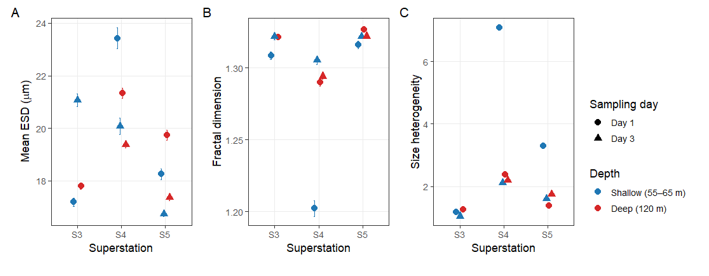

**Supplementary Figure 2.** Particle morphological characteristics across stations (S3-S5), sampling days (Day 1 and Day 3), and depths (shallow: 55-65 m; deep: 120 m). A) mean equivalent spherical diameter (ESD), B) mean fractal dimension, and C) size heterogeneity (coefficient of variation of particle area). Point colour indicates sampling depth and symbol indicates sampling day.

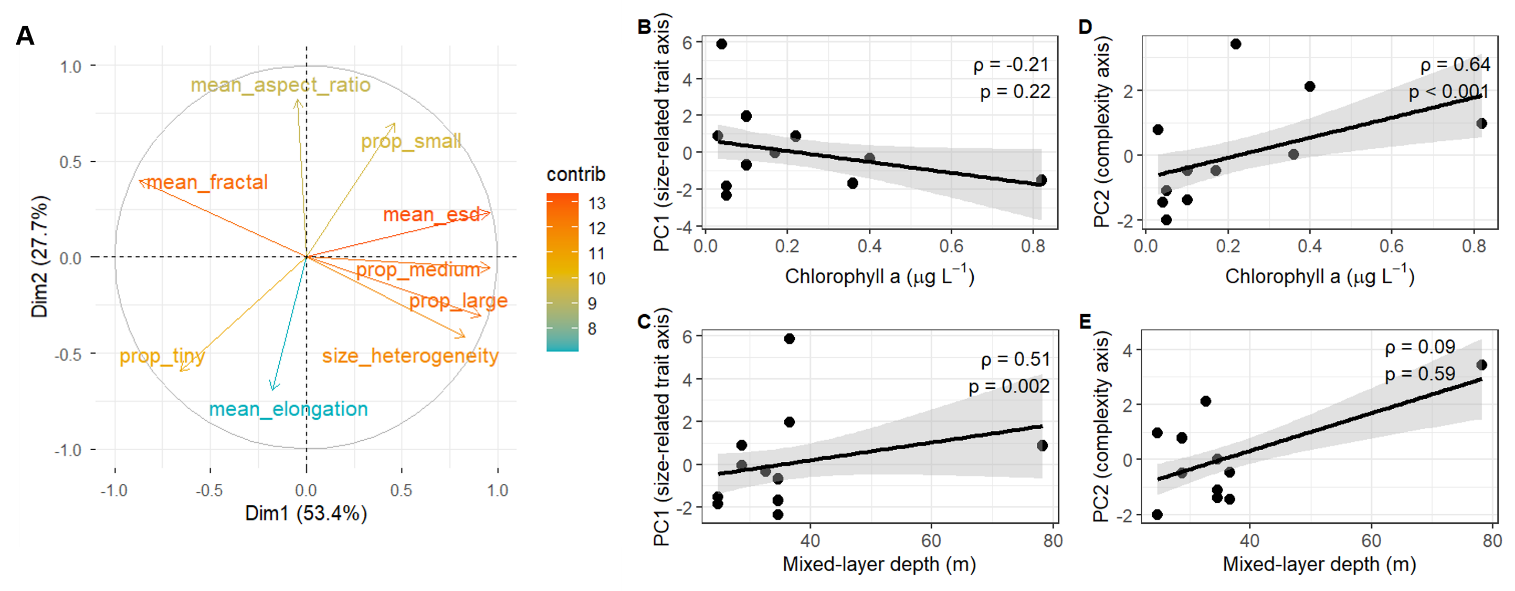

**Supplementary Figure 3.** A) Principal component analysis (PCA) biplot illustrating loadings of particle morphological variables on the first two principal components. PC1 (Dim 1; 53.4% of variance) represents a particle size-related trait axis associated with equivalent spherical diameter (ESD), particle size-class composition and size heterogeneity, while PC2 (Dim 2; 27.7%) captures variation in particle geometry and structural complexity associated with fractal dimension, elongation and aspect ratio. Vectors are coloured by variable contribution to each axis. B) Relationship between chlorophyll a concentration and the particle size-related trait axis (PC1). C) Relationship between mixed layer depth and the particle size-related trait axis (PC1). D) Relationship between chlorophyll a concentration and the complexity-related trait axis (PC2). E) Relationship between mixed layer depth and the complexity-related trait axis (PC2). Black lines indicate fitted linear regressions and grey shading represents 95% confidence intervals.

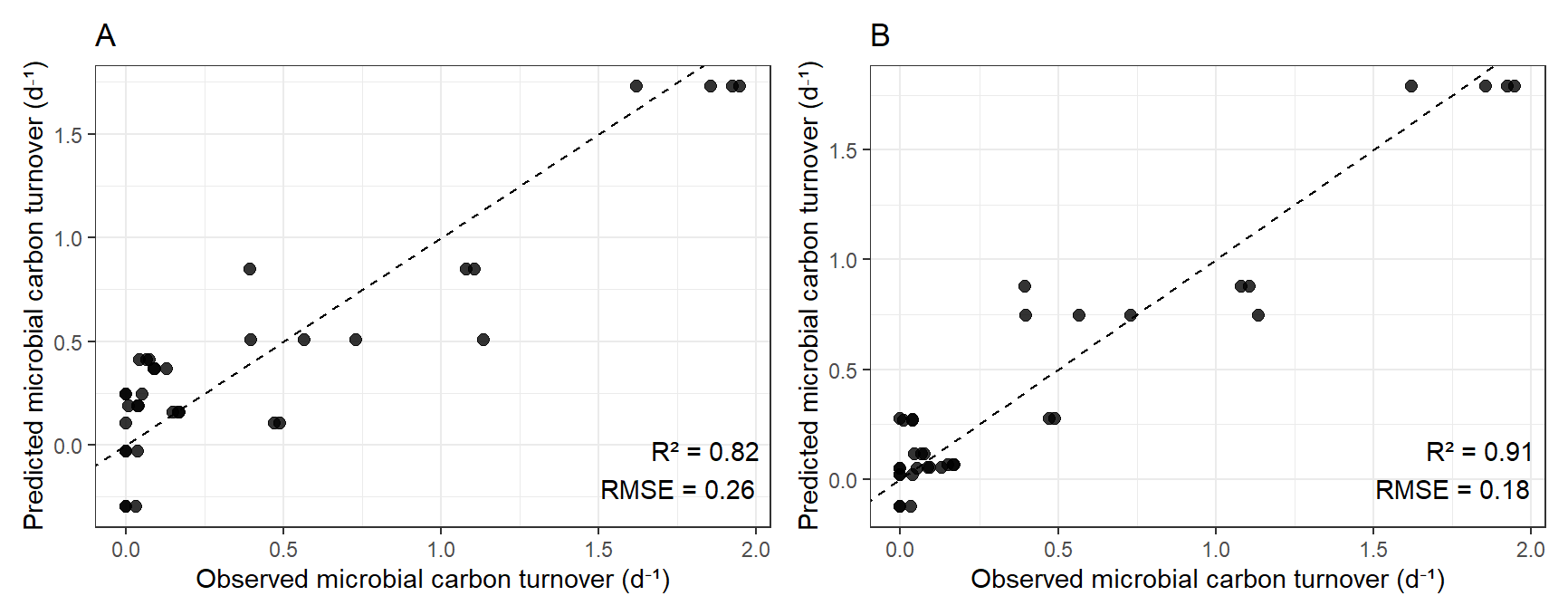
**Supplementary Figure 4.** Predicted vs observed microbial carbon turnover for A) environmental and B) particle informed (environment + particle traits) linear models. The environmental model includes chlorophyll a, dissolved oxygen and mixed layer depth. The particle informed model includes the environmental model parameters and particle traits of size and complexity. Dashed lines indicate 1:1 agreement between observed and predicted values. Points represent individual incubations. Root mean square error (RMSE).

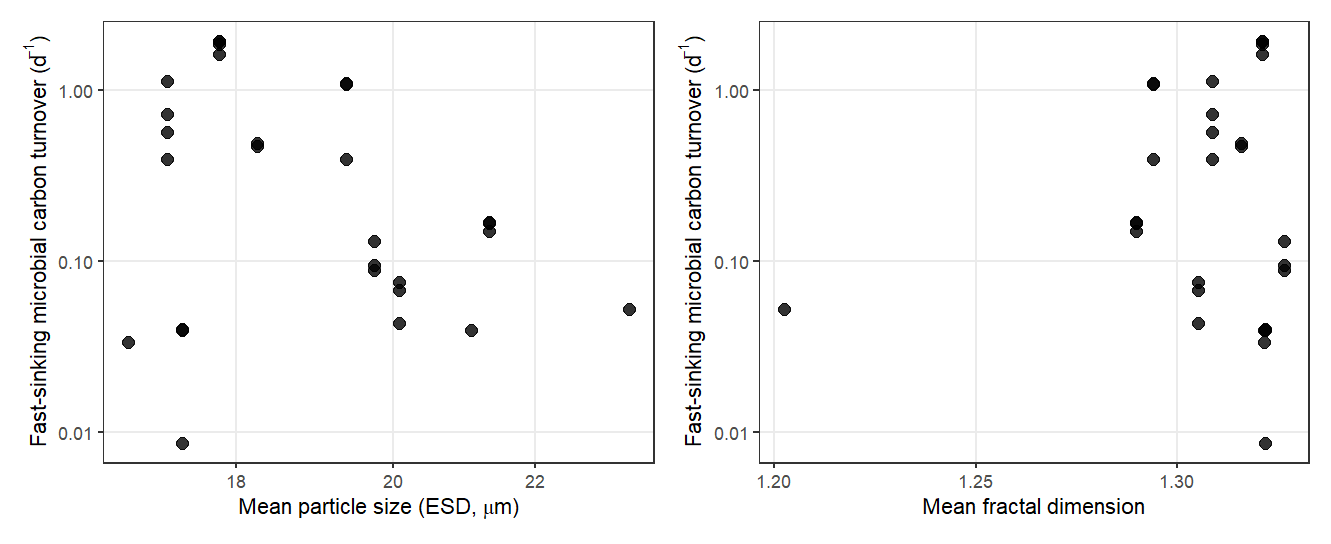

**Supplementary Figure 5.** Plots of A) mean particle equivalent spherical diameter (ESD) and B) mean fractal dimension against microbial carbon turnover (d^-1^).

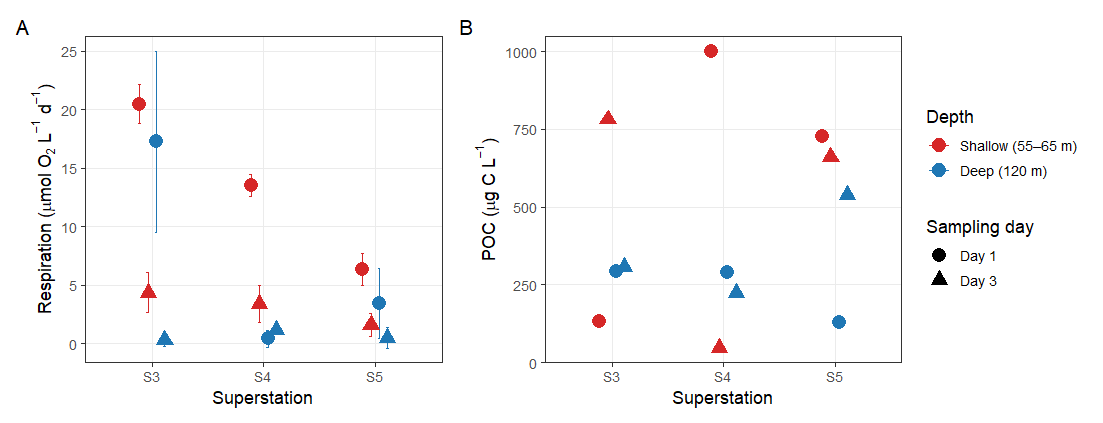

**Supplementary Figure 6.** Respiration and particulate organic carbon (POC) associated with fast-sinking particles across superstations (S3-S5), sampling days (Day 1 and Day 3), and depths. **(A)** Mean respiration rates (± SD) of replicate incubations, expressed as μmol O₂ L⁻¹ d⁻¹. **(B)** Particulate organic carbon concentrations in incubation vials (μg C L⁻¹). Red symbols indicate shallow samples (55-65 m) and blue symbols indicate deep samples (120 m). Circles represent Day 1 samples and triangles represent Day 3 samples.

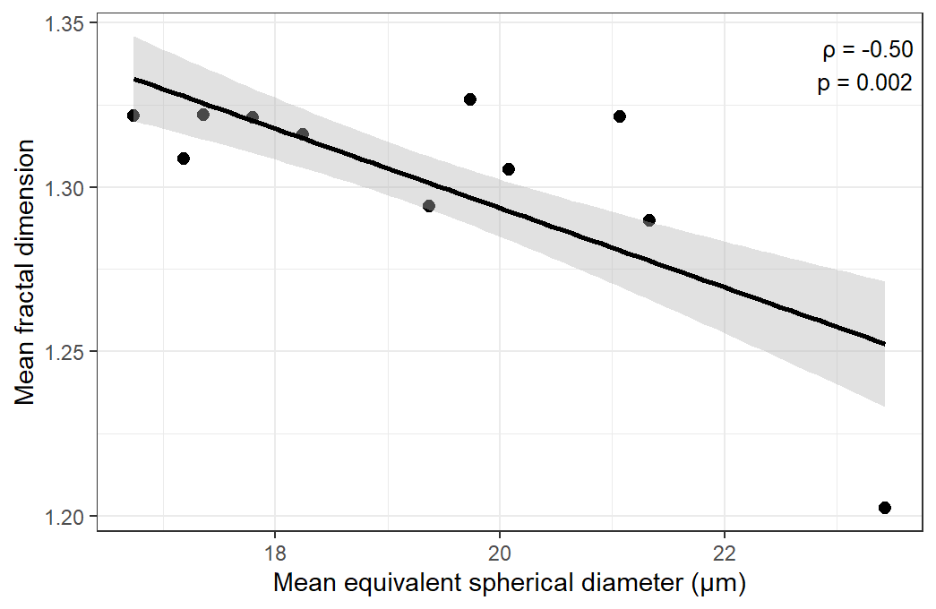

**Supplementary Figure 7.** Relationship between mean equivalent spherical diameter (ESD) and mean fractal dimension across particle assemblages. Structural complexity decreased with increasing particle size (Spearman's ρ = −0.50, p = 0.002). Black line indicates fitted linear regression and grey shading represents the 95% confidence interval.

**Supplementary Tables**

**Supplementary Table 1.** Sampling depths, times and associated CTD deployment and mixed layer depths for microbial carbon turnover incubations. Mixed Layer Depth (MLD) was determined from the CTD cast closest in time to each MSC deployment using a density threshold criterion of Δσθ = 0.05 kg m⁻³.

| **Date** | **Station** | **Day** | **Shallow Depth (m)** | **Time of shallow MSC firing** | **Deep Depth**  **(m)** | **Time of deep MSC firing** | **Time of nearest CTD deployment** | **Mixed Layer Depth (m)** | **Euphotic Depth (m)** |
| --- | --- | --- | --- | --- | --- | --- | --- | --- | --- |
| 06-06-2026 | S3 | 1 | 65 | 06:47 | 120 | 07:01 | 04:21 | 25 | 43 |
| 08-06-2026 | S3 | 3 | 65 | 03:58 | 120 | 04:19 | 02:29 | 78 | 43 |
| 12-06-2026 | S4 | 1 | 55 | 06:43 | 120 | 07:14 | 04:40 | 37 | 31 |
| 14-06-2026 | S4 | 3 | 55 | 04:08 | 120 | 04:25 | 02:33 | 29 | 15 |
| 17-06-2026 | S5 | 1 | 55 | 09:02 | 120 | 09:06 | 07:59 | 33 | 25 |
| 19-06-2026 | S5 | 3 | 55 | 06:03 | 120 | 06:21 | 04:59 | 35 | 15 |

**Supplementary Table 2.** Summary table of parameter estimates of the particle trait-informed linear model (environmental parameters + particle traits) of microbial carbon turnover. Coefficients are reported with standard error (SE), t-values, and p-values.

| **Predictor** | **Environmental model estimate** | **SE** | **t-value** | **p-value** | **Particle-trait model estimate** | **SE** | **t-value** | **p-value** |
| --- | --- | --- | --- | --- | --- | --- | --- | --- |
| Intercept | 23.3 | 3.66 | 6.38 | <0.001 | 26.15 | 2.83 | 9.23 | <0.001 |
| Chlorophyll-*a* | 3.54 | 0.367 | 9.65 | <0.001 | 4.19 | 0.30 | 13.76 | <0.001 |
| Oxygen | -0.084 | 0.013 | -6.22 | <0.001 | -0.096 | 0.010 | -9.23 | <0.001 |
| Mixed-layer depth | -0.017 | 0.003 | -5.09 | <0.001 | -0.004 | 0.004 | -1.08 | 0.29 |
| PC1 (Size) | - | - | - | - | -0.065 | 0.018 | -3.66 | 0.001 |
| PC2 (Complexity) | - | - | - | - | -0.144 | 0.038 | -3.83 | <0.001 |
